# *De novo* design of stable proteins that efficaciously inhibit oncogenic G proteins

**DOI:** 10.1101/2023.03.28.534629

**Authors:** Matthew C. Cummins, Ashutosh Tripathy, John Sondek, Brian Kuhlman

## Abstract

Many protein therapeutics are competitive inhibitors that function by binding to endogenous proteins and preventing them from interacting with native partners. One effective strategy for engineering competitive inhibitors is to graft structural motifs from a native partner into a host protein. Here, we develop and experimentally test a computational protocol for embedding binding motifs in de novo designed proteins. The protocol uses an “inside-out” approach: Starting with a structural model of the binding motif docked against the target protein, the de novo protein is built by growing new structural elements off the termini of the binding motif. During backbone assembly, a score function favors backbones that introduce new tertiary contacts within the designed protein and do not introduce clashes with the target binding partner. Final sequences are designed and optimized using the molecular modeling program Rosetta. To test our protocol, we designed small helical proteins to inhibit the interaction between Gα_q_ and its effector PLC-β isozymes. Several of the designed proteins remain folded above 90°C and bind to Gα_q_ with equilibrium dissociation constants tighter than 80 nM. In cellular assays with oncogenic variants of Gα_q_, the designed proteins inhibit activation of PLC-β isozymes and Dbl-family RhoGEFs. Our results demonstrate that computational protein design, in combination with motif grafting, can be used to directly generate potent inhibitors without further optimization via high throughput screening or selection.

**statement for broader audience:** Engineered proteins that bind to specific target proteins are useful as research reagents, diagnostics, and therapeutics. We used computational protein design to engineer de novo proteins that bind and competitively inhibit the G protein, Gα_q_, which is an oncogene for uveal melanomas. This computational method is a general approach that should be useful for designing competitive inhibitors against other proteins of interest.

## Introduction

One strategy for imparting function to designed proteins is to embed preexisting binding motifs within the designs. This strategy has been used to create new inhibitors, metal-binding proteins, and immunogens for subunit vaccines^1–5^. However, several design requirements should be met when embedding a new binding motif within a protein to create a well-behaved, high-affinity binder. For example, the host protein should stabilize the binding residues in a binding-competent conformation, the binding surface should be accessible to the target molecule, and the host protein itself should be well-folded and soluble. One effective approach for meeting these design requirements is to use naturally occurring proteins as the host scaffold. Host proteins can be identified by scanning the protein structural database (PDB) for proteins that contain structural elements that geometrically match the binding motif of interest^6^. However, a major limitation of relying on naturally occurring proteins is that as binding motifs become larger and more complicated, it is often challenging to find a naturally occurring protein that is a good match.

An alternative strategy for scaffolding binding motifs is to embed them in de novo designed proteins specifically created to present the motif of interest. There has been considerable progress in the field of de novo protein design as methods have been developed for constructing protein backbones that satisfy predetermined geometric constraints and are inherently designable, i.e., the residues are appropriately positioned relative to each other to allow favorable packing interactions between side chains. These backbone generation approaches include parametric specification of common structural motifs (e.g., coiled coils)^7, 8^, backbone assembly from pieces of naturally occurring proteins^9–12^, and most recently, the training of neural networks to generate novel protein backbones^13–17^. Here, we test whether the backbone generation protocol SEWING (Structure Extension WIth Native fragment Graphs) can create protein binders by embedding a protein binding motif in de novo designed proteins^18^. SEWING assembles backbones in a stepwise fashion from naturally occurring helix-turn-helix (HTH) motifs taken from the PDB^19^. During SEWING, HTH motifs are combined by superimposing the C-terminal helix of one HTH motif with the N-terminal helix of another HTH motif. As the protein grows, a low-resolution score function is used to ensure that no backbone clashes are introduced, and that helices are placed at favorable distances relative to each other. To incorporate a binding motif in a backbone generated with SEWING, the binding motif is used as the starting point for backbone assembly.

As a test case for designing protein binders with SEWING, we designed and characterized proteins that competitively bind the G protein Gα_q_ at its natural binding site for phospholipase C-β isozymes (PLC-β). The G protein Gα_q_ binds and activates phospholipase C-β (PLC-β) isozymes for the catalysis of phosphatidylinositol 4,5-bisphosphate (PIP_2_) into diacylglycerol (DAG) and inositol 1,4,5-trisphosphate (IP_3_) ^20, 21^. Activation of PLC-β isozymes by Gα_q_ is essential for various cellular processes linked to proliferation and differentiation^22^. Furthermore, mutations that constitutively activate Gα_q_ and promote its binding to PLC-β drive ∼90% of all uveal melanomas^23, 24^. Recently, new functions of Gα_q_ have been identified using pharmacological tools that selectively turn off Gα_q_ signaling ^25–27^. The most common tool is the naturally occurring cyclic depsipeptide FR900359^28^. Unfortunately, FR is challenging to synthesize, and the purification from its natural source is laborious^29^. Genetically encoded inhibitors specific for Gα_q_ are likely to serve as useful research reagents for studying G protein signaling.

The interaction between Gα_q_ and PLC-β3 is mediated by a helix-turn-helix (HTH) motif from PLC-β3 that docks against the surface of Gα_q_^30^. Mutating residues at this interface disrupts the activation of PLC-β3 and an affinity-optimized peptide derived from the HTH binds to Gα_q_ with an equilibrium dissociation constant (K_D_) of 400 nM. In a previous study, we explored if binding could be tightened further by embedding the PLC-β3 HTH in a naturally occurring folded protein to stabilize the peptide in a binding-competent conformation^31^. After several rounds of yeast cell surface display with a protein library containing over 1 million sequence variations, we identified an engineered protein, BB25, that binds the active form of Gα_q_ with a K_D_ of 20 nM. This result demonstrated the usefulness of embedding binding motifs in a folded protein but required extensive screening and selection. In this current study, we address whether de novo protein design can achieve a similar outcome without constructing a large protein library.

## Results

### SEWING backbones stabilize the HTH

The crystal structure of PLC-β3 bound to Gα_q_ (PDB entry 7SQ2) was used as the starting point for protein design simulations (Figure 1). All residues from PLC-β3 except for the HTH (residues 852-875) were removed from the structure, and SEWING was used to build a protein backbone around the HTH. The building blocks for SEWING were thousands of HTH fragments extracted from the PDB. During SEWING, new HTH fragments are added to the C-terminus of a growing polypeptide chain by superimposing the N-terminal helix of a HTH fragment with the C-terminal helix of the design model. The reverse superposition is performed to grow the N-terminus of the design model. A Monte Carlo procedure is used to optimize the score of the design models by swapping in and out alternative HTH fragments and by sampling alternative alignments during superposition of the helices. For putative binders of Gα_q_, the primary score term used during backbone assembly was Rosetta’s motif score which is independent of amino acid sequence, and rewards helices placed near each other at the appropriate distance to allow for favorable side chain packing between smaller hydrophobic amino acids (Ile, Leu, Val, Ala). A clash term was also used to ensure that helices did not overlap or clash with Gα_q_.

**Figure 1.**
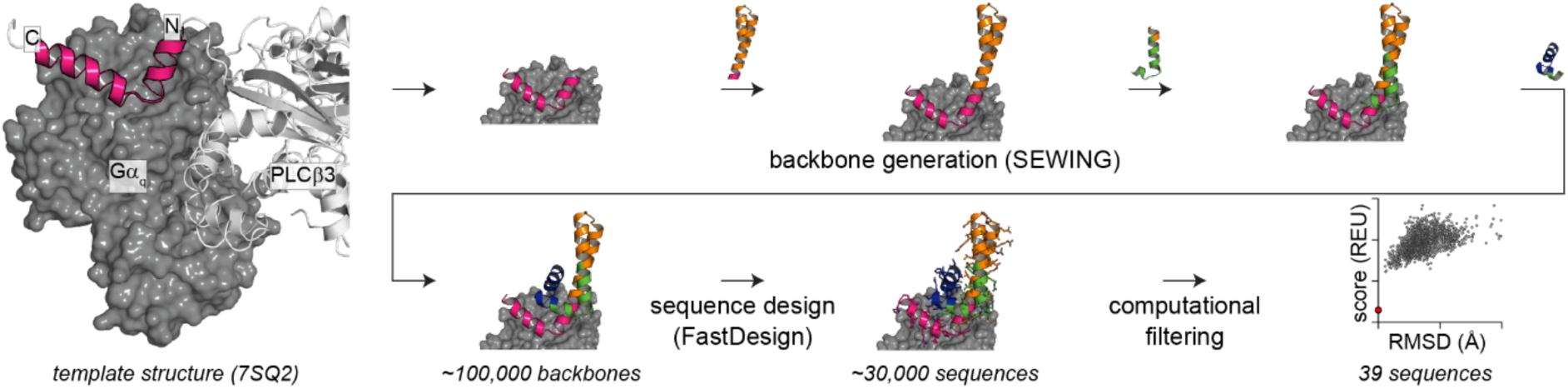
SEWING builds de novo backbones by combining helix-turn-helix (HTH) fragments extracted from the PDB. To design Gα_q_ inhibitors with SEWING, the three-dimensional coordinates of Gα_q_ (dark gray) along with residues 852-875 from PLC-β3 (magenta) were taken from a crystal structure of PLC-β3 bound to Gα_q_ (PDB code 7SQ2). SEWING was then used to “grow” a protein around the PLC-β3 motif by adding new helices to either terminus of the protein. After backbone assembly, a sequence was designed for the de novo protein using Rosetta’s FastDesign protocol. The top scoring sequences were filtered using scoring metrics and selected for experimental validation.

Two alternative SEWING protocols were used to create two families of de novo designs. Protocol-1 used the standard Rosetta motif score to evaluate packing between helices, while protocol-2 adjusted the motif score so that interactions between residues distant in primary sequence were upweighted relative to interactions close in primary sequence. This change was made to favor higher contact order structures that contain interactions between the protein’s N- and C-terminal regions. Protocol-1 was used to create proteins with 4 helices, while protocol-2 was used to create proteins with 5 or 6 helices. After the protein backbones were generated with one of the two SEWING protocols, the FastDesign method in Rosetta was used to design sequences for each model^32^. FastDesign iterates between rotamer-based sequence design and backbone refinement to find low-energy sequence/structure pairs. Gα_q_ was present in the models during FastDesign, and PLC-β3 HTH residues that make contact with Gα_q_ were not allowed to mutate during FastDesign. Residues in the rest of the design model were allowed to mutate to any residue except cysteine.

The models produced with FastDesign were ranked using the Rosetta full-atom energy function, and the lowest ∼100 scoring sequences were evaluated with Rosetta’s ab initio structure prediction protocol. Sequences that had a high prediction to fold to the design model (probability > 80%) were selected for experimental studies (17 sequences from protocol-1 and 22 sequences from protocol-2). The proteins were expressed in E. coli and purified using metal affinity chromatography. Eleven of the 17 protocol-1 designs and 10 of the 22 protocol-2 designs expressed well and were soluble after affinity purification. Preliminary binding measurements were performed using biolayer interferometry to identify which designs bind most tightly to Gα_q_. The three tightest binders from protocol-1 (SEWN1.7, SEWN1.9, and SEWN1.12) and protocol-2 (SEWN2.12, SEWN2.16, and SEWN2.20) were selected for more extensive biophysical characterization.

### SEWING designs are stable and bind Gα_q_ with nanomolar affinity

Per design specifications, the SEWN2.0 sequences are longer than the SEWN1.0 sequences, and more contacts are made between the N- and C-termini in the SEWN2.0 design models (Figure 2A). Far-UV circular dichroism (CD) spectra at 20°C of all 6 designs exhibit minima at 222 nm and 208 nm, a feature characteristic of helical proteins (Figure 2B). For the designs SEWN1.7 and SEWN1.9, the CD signal is more negative at 208 nm than 222 nm, indicating the proteins are less helical than the other four designs. The lower helicity for SEWN1.7 and SEWN1.9 could indicate that a population of the proteins is unfolded at room temperature or that only a portion of the protein adopts a helical conformation. The stability of all 6 designs was probed by monitoring the CD signal at 222 nm as a function of temperature (Figure 2C) or concentration of guanidine hydrochloride (GuHCl) (Figure 2D). In the temperature melts, all the proteins lose helicity as the temperature is raised, but the CD signal at 222 nm remains more negative at high temperatures for the SEWN2.0 designs. Consistent with SEWN1.7 and SEWN1.9 being partially unfolded at room temperature, the CD signal for these proteins decreases as soon as low amounts of chemical denaturant are added. In contrast, SEWN 1.12 and SEWN 2.16 do not appreciably unfold until the addition of 3 M and 6 M GuHCl, respectively, while SEWN 2.12 and 2.20 resist unfolding at all concentrations of GuHCl.

**Figure 2.**
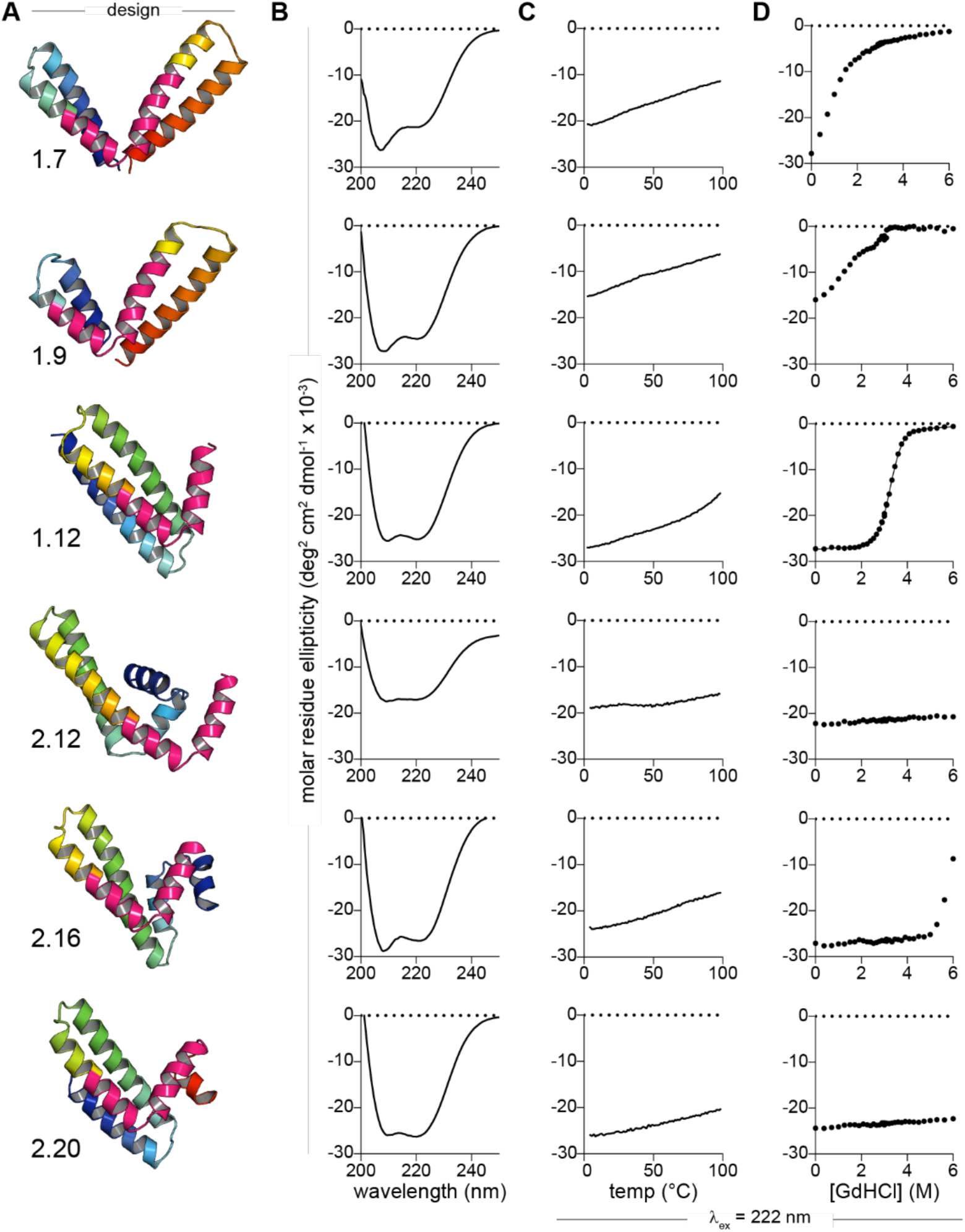
Secondary structure and stability of SEWN proteins. **A.** Rosetta models of the SEWN proteins with the HTH motif from PLC-β3 highlighted in hot pink. **B.** Circular dichroism (CD) spectra of the SEWN proteins display negative peaks at 208 and 222 nm, consistent with the formation of α-helices. **C.** CD signal at 222 nm as a function of temperature. SEWN1.12 and all of the SEWN2 proteins maintain large negative ellipticity at high temperatures. **D.** CD signal at 222 nm as a function of guanidine hydrochloride concentration.

To probe the oligomerization state of the 6 de novo designs, size exclusion chromatography with multi-angle light scattering (SEC-MALS) was performed (Figure S1). At protein concentrations of ∼20 μM, SEWN1.7, SEWN1.12, SEWN2.12 and SEWN2.16 are all monomeric. At these concentrations, the design SEWN1.9 appears to exist as a mixture of monomers and oligomers, and the design SEWN2.20 is primarily a tetramer.

The binding affinities between the designs and Gα_q_ were measured with both biolayer interferometry (BLI) and a competitive binding assay based on fluorescence anisotropy measurements. In the BLI experiments, biotin-labeled Gα_q_ (labeled via an AviTag) was immobilized on streptavidin biosensors and incubated with various concentrations of the de novo designed proteins. The measured equilibrium dissociation constants (K_D_) varied from 83 nM (SEWN2.20) to less than 1 nM (SEWN1.7) (Figure 3A). The tight affinity exhibited by SEWN1.7 reflects a very slow dissociation rate, <1.0×10^−7^ sec^−1^ (Table S1). These BLI experiments were repeated but in the absence of AlF_4_^−^ (Figure S2). The SEWN proteins show over 15 fold less response at twice the concentration of SEWN protein. The BLI experiments indicate that all six designs bind tightly to the active form of Gα_q_, but they do not indicate where the designs are binding on Gα_q_.

**Figure 3.**
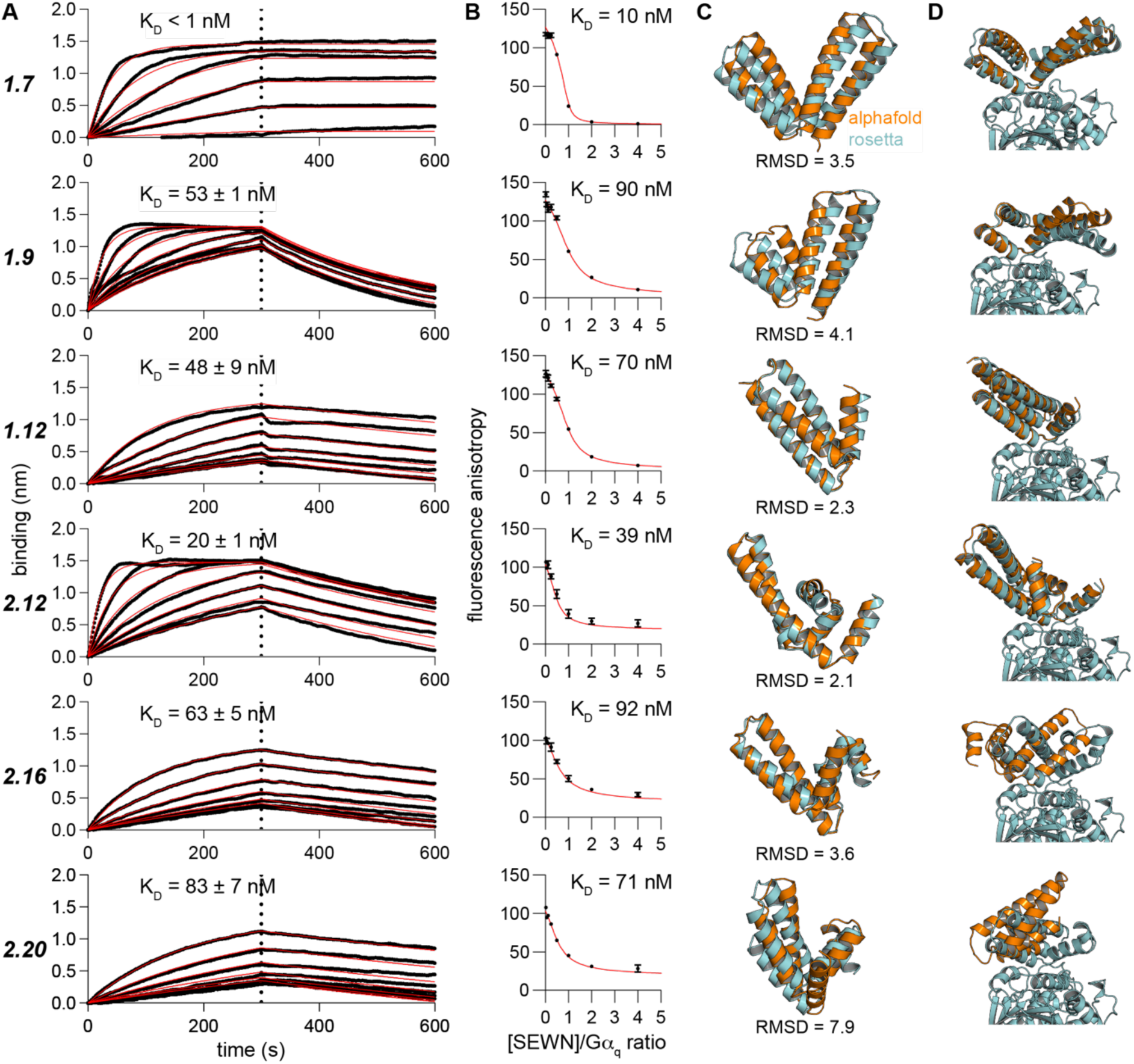
The SEWN proteins bind Gα_q_ at the predicted binding site. **A.** The affinities of the designed proteins for Gα_q_ were measured with biolayer interferometry (BLI). Biotinylated Gα_q_ was immobilized on streptavidin sensors and the sensors were dipped into solutions of the SEWN proteins at concentrations varying between 0 nM and 500 nM. **B.** Competitive fluorescence anisotropy experiments corroborate the BLI. Gα_q_ (1 μM) was preincubated with 20 nM FAM-labeled HTH peptide, and anisotropy was measured as a function of the concentration of SEWN protein. The fitted curves are from a mathematical model of competitive binding that adjusts the K_d_ as well as the starting and ending anisotropy values to optimize the fit.^31^ **C.** AlphaFold2 predictions of the SEWN proteins with the Cα RMSD indicated for each design. **D.** AlphaFold2 predictions of the SEWN proteins as a complex with Gα_q_. The structures were aligned by superimposing the Gα_q_ chains only. The Gα_q_ chain from the AlphaFold2 predictions are hidden for clarity.

To probe if the designs are binding to the target binding site on Gα_q_ and if they can compete with the binding of PLC-β isozymes, competitive binding experiments were performed with an affinity-optimized HTH peptide derived from PLC-β3 (Figure 3B). In these experiments, the HTH peptide conjugated to FAM was pre-mixed with Gα_q_ and the designed protein was titrated into the sample. If binding is competitive, then the designed protein should displace the peptide leading to a decrease in fluorescence anisotropy. All six designs efficiently displaced the HTH peptide, which has a K_D_ of 400 nM for Gα_q_. The competitive binding curves were fit with a mathematical model to determine the K_D_ of the designs for Gα_q_. The fitted K_D_s agreed well with the BLI measurements, varying less than 2-fold from K_D_s determined by BLI. These experiments confirm that the designed proteins bind to the effector site on Gα_q_ used to engage PLC-β isozymes.

### AlphaFold2 models agree with some Rosetta-predicted models of designs and complexes

The computational models produced in this study were all created before the public release of the structure prediction software, AlphaFold2. AlphaFold2 has shown excellent performance in the Critical Assessment of Structure Prediction (CASP) and provides an orthogonal method for validating designed proteins. Our goal was to design de novo proteins that stabilize the HTH motif in a binding competent conformation. Therefore, we first examined AlphaFold2 predictions for the design sequences without Gα_q_ (Figure 3C). The overall topologies and secondary-structure pattern are highly similar in the AlphaFold2 and Rosetta models. Structural differences indicative of hinge motion around the HTH motif in SEWN1,7 and SEWN2.12 suggest there may be flexibility in this region of the proteins. The Rosetta models for SEWN1.12 and SEWN2.12 most closely match the AlphaFold2 predictions with a Cα RMSD of 2.3 and 2.1 Å, respectively.

To evaluate the designed interfaces, we used AlphaFold2 to predict the structure of the complexes formed between the SEWN designs and Gα_q_ (Figure 3D). Once again, we measured the Cα RMSD of the AlphaFold2 predictions to the Rosetta models, but we first superimposed the Gα_q_ chains so that the measurement reflects both changes in backbone conformation and differences in the positioning of the designs against Gα_q_. The lowest RMSDs were observed for SEWN1.12 (2.2 Å) and SEWN2.12 (2.1 Å) (Table S2). Unlike the other four designs, the AlphaFold2 models for SEWN2.16 and SEWN2.20 do not place the HTH motif in the binding groove of Gα_q_. As both designs bind to Gα_q_ with K_D_s below 100 nM, it is likely that the AlphaFold2 predictions are not accurately modeling the structures of the bound complexes for these cases.

### SEWING designs inhibit Gα_q_ in HEK293 cells

Two cellular activity assays were performed to determine if SEWN1.12 and SEWN2.12 can inhibit Gα_q_ signaling in cells. SEWN1.12 and SEWN2.12 were selected for cellular assays because the proteins are well-folded, monomeric, bind to Gα_q_ with K_D_s below 50 nM, and there is a close match between the AlphaFold2 and Rosetta models for these designs. First, we tested whether the designs inhibit Gα_q_-mediated activation of PLC-β isozymes in HEK293 cells. In this assay, a constitutively active mutant of Gα_q_(Q209L) implicated in uveal melanoma is expressed in the cells, which activates PLC-β isozymes and leads to the production of [^3^H]inositol phosphates that are detected using a scintillation counter. To test for inhibition of Gα_q_ by the designed proteins, the designs were co-transfected along with Gα_q_(Q209L), and inositol phosphate levels were measured. As Gα_q_ and PLC-β are active at the plasma membrane, two types of constructs were tested. The designed inhibitors were either expressed as cytosolic proteins or as fusions with a C-terminal CAAX motif for localization to the plasma membrane. The affinity-optimized HTH peptide from PLC-β was also encoded for mammalian expression and tested in the inhibition assays. All of the proteins expressed at similar levels inside the cells as judged by western blotting (Figure 4A). When expressed without a membrane anchor, SEWN1.12 and SEWN2.12 were significantly more efficacious than the HTH alone (P < 0.0001) and produced a ∼4-fold drop in inositol phosphate levels. When expressed with CAAX-box, all three inhibitors performed similarly with inhibition levels comparable to those observed with the cytosolic expressed SEWN1.12 and SEWN2.12. These results indicate that some feature of the SEWN designs, perhaps their higher affinity for Gα_q_ or the fact that they are folded proteins, precludes them from needing a membrane localization tag to inhibit Gα_q_ efficaciously. We also tested an additional variant of Gα_q_ (Q209P) in the inhibition assay that is constitutively active and observed in uveal melanoma (Figure S3). The results with Gα_q_ Q209P closely mirror those observed with Gα_q_ Q209L.

**Figure 4.**
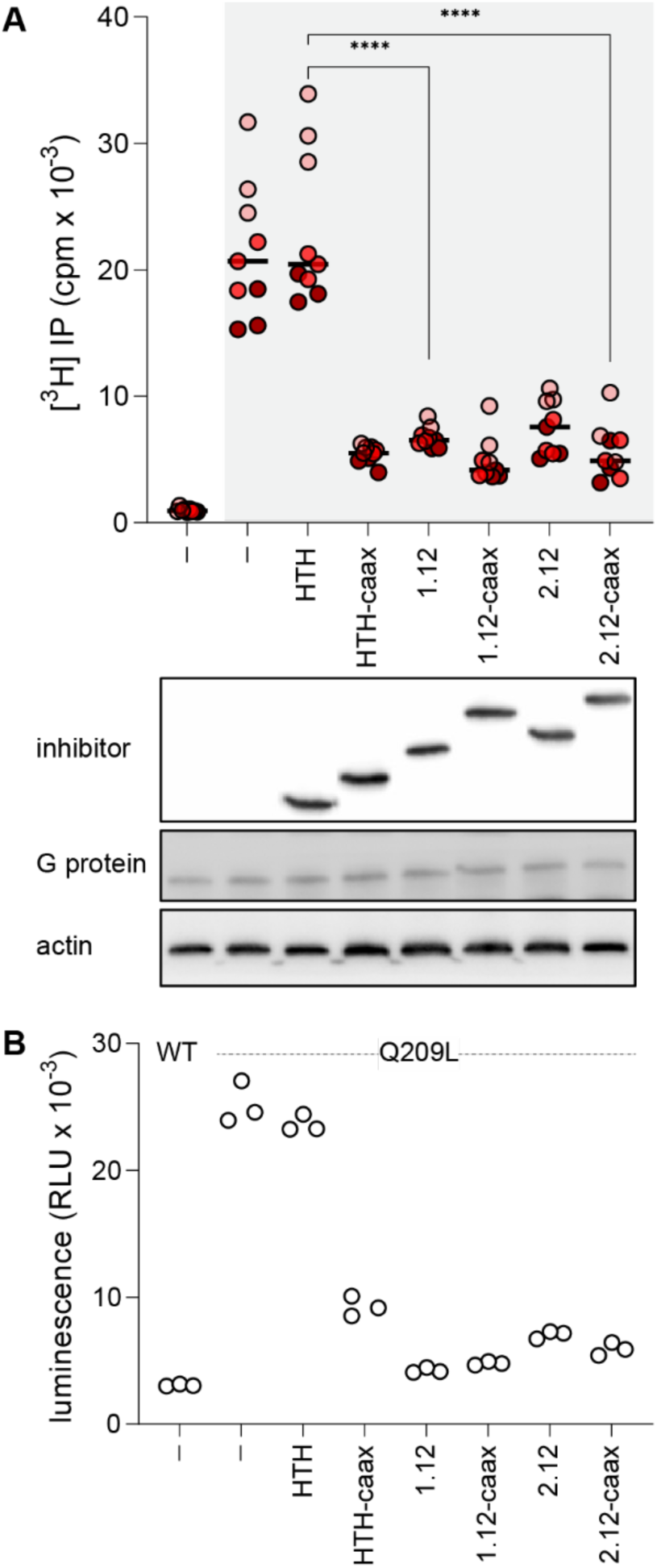
SEWN1.12 and SEWN2.12 efficaciously inhibit Gα_q_ signaling in HEK293 cells. **A.** SEWN1.12 and SEWN2.12 inhibit Gα_q_-mediated activation of PLC-β isozymes. Columns in white were cells that were transfected with WT Gα_q_. Columns with gray background were cells that were transfected with Q209L Gα_q_. Inositol phosphates produced from HEK293 cells were measured after being transfected with various constructs. SEWN1.12 and SEWN2.12 are significantly more efficaciously than HTH alone. However, upon localization to the membrane via a CAAX tag, all constructs inhibit Gα_q_ with similar degrees of inhibition. The experiment was performed 3 times. Each circle represents a unique well of cells that was individually transfected and measured. Circles that are colored the same represent replicates that were done using cells that were split from the same HEK293 culture. **B.** The SEWN proteins also inhibit Gα_q_-mediated activation of Dbl family RhoGEFs. The luciferase activity of HEK293 cells were measured after transfecting them with a luciferase under an SRE promoter, along with various inhibitory constructs. Similar to the PLC-β isozyme inhibition, SEWN1.12 and SEWN2.12 efficaciously inhibit activation of Dbl family RhoGEFs.

Next, we tested if the designs inhibit other effectors of Gα_q_ (Figure 4B). Gα_q_ activates Dbl-family RhoGEFs by engaging HTH motifs structurally similar to the HTH on PLC-β3. Furthermore, the HTH from Dbl-family RhoGEFs binds Gα_q_ on the same surface as the HTH from PLC-β3. Thus, if the designs can inhibit the activation of PLC-β isozymes, they should also inhibit the activation of Dbl-family RhoGEFs. We tested the activation of Dbl-family RhoGEFs by transfecting HEK293 cells with plasmids encoding firefly luciferase under an SRE promoter. If the RhoGEFs are active, the SRE transcription factor will be active, and the luciferase will be expressed. Like the inhibition assay with PLC-β isozymes, SEWN1.12 and SEWN2.12 efficaciously inhibit Gα_q_ without a CAAX tag, while the HTH peptide does not inhibit Gα_q_ without the addition of a CAAX tag.

## Discussion

Our results demonstrate that SEWING can be used to embed structural motifs in stable de novo designed proteins to maintain or enhance the starting motif’s activity. All six designs that were extensively characterized bind more tightly to Gα_q_ than the affinity-optimized HTH peptide they were derived from. It is interesting to consider why there is an increase in binding affinity. In the design models and the AlphaFold predictions of the SEWN / Gα_q_ complexes, the interface is formed almost entirely from interactions between the HTH motif and Gα_q_, with the design process adding few additional contacts. Since new contacts are unlikely to be the source of increased affinity, an alternative explanation is that the design process has pre-stabilized the HTH motif in a binding competent conformation and reduced conformational entropy losses associated with binding. We previously demonstrated that the HTH peptide adopts a random coil in the unbound state^33^, and thus undergoes significant structural ordering upon binding.

The stability of the six designs was highly varied. SEWN 1.7 and 1.9 begin to unfold immediately when chemical denaturant is added, while 2.12 and 2.20 remain fully helical up to 6 M guanidine hydrochloride. These differences likely reflect the tertiary folds of the designs. SEWN 1.7 and 1.9 are predicted to adopt a “rabbit ear” like structure with few contacts between the N- and C-terminus and the HTH motif acting as a hinge between the two halves of the protein. The more stable designs were created with protocol 2, which biased the SEWING assembly process to form contacts between the N- and C-terminus of the molecule and resulted in tertiary folds that are more globular. It has been observed that proteins from thermophiles tend to have higher contact order than their mesophilic homologs^34^.

SEWN 1.7 exhibits the tightest affinity for Gα_q_ despite only being moderately stable. It may be that the more stable designs present the HTH motif in a conformation that does not perfectly match the bound conformation, and therefore there is an energetic cost for rearranging the binding motif upon binding. In contrast, SEWN 1.7 is likely to be more flexible about the HTH hinge, and therefore there may be a smaller energetic cost for any conformational changes needed for binding. Another potential explanation is that the angle between the first and second helix of the HTH may adopt a range of values and still bind Gα_q_. Thus, stabilizing each helix while allowing a range of angles between the helices results in tighter affinity.

In a previous study, we examined if high-affinity binders for Gα_q_ could be engineered by embedding the HTH from PLC-β3 in a naturally occurring well-folded protein. A high-affinity binder was generated, but several limitations of the approach became apparent during the project. For example, only two proteins from the entire PDB were identified that could appropriately present the HTH motif, and it is likely that if the motif were more complicated or discontinuous, no matches would have been identified. Here, by using de novo protein design, we were not limited to structures previously deposited in the PDB. Furthermore, instead of using a natural protein that was evolved for some unrelated function, these de novo proteins were explicitly designed to be well-folded and to bind and competitively inhibit Gα_q_. In the previous study, it was also necessary to screen a library >1×10^7^ protein variants to identify binders for Gα_q_ with equilibrium dissociation constants equivalent to those reported here.

One motivation for this study was the prediction that a tighter binder would have more efficacious inhibition of Gα_q_. Indeed, both SEWN1.12, and SEWN2.12 bind Gα_q_ with a K_D_ that is five times tighter than the HTH alone and are significantly more efficacious than the unanchored HTH in HEK293 cells. However, once a CAAX tag is added for membrane localization, all of the constructs perform similarly. The ability to inhibit Gα_q_ without the use of a CAAX tag is potentially valuable for experiments where the goal is to turn off Gα_q_ signaling throughout the cell. For example, Gα_q_ is known to signal in early endosomes^35–38^. One current hypothesis is that early endosome signaling allows for sustained G protein signaling, since plasma membrane signaling is dampened by β-arrestins. However, this hypothesis and many others require further investigation. Thus, having an inhibitor that can globally turn off Gα_q_ signaling or be specifically sequestered in endosomes or the plasma membrane is a valuable feature for dissecting the roles of Gα_q_ signaling from its various locations.

Very recently, there has been exciting progress in using deep learning to train neural networks that can predict protein structure and generate novel backbones that are good templates for protein design. Like SEWING, these methods can scaffold binding motifs in a manner that maintains or enhances functionality^13, 14^. One advantage of these approaches over SEWING is that they are compatible with all types of protein folds (α, β, mixed α/β) and can readily generate large protein structures (> 300 residues). As methodologies continue to advance, it will be exciting to see if multiple functions can be incorporated into single proteins to create signaling systems that compare in complexity to naturally occurring systems.

## Materials and methods

### Computational protein design using Rosetta

For protocol 1 designs, PDB 7SQ2 was used as the structure of HTH bound to Gα_q_. Residues 852-875 from PLC-β3 were used as the starting HTH on which SEWING appended helical substructures. Approximately 50,000 SEWING assemblies were made. SEWING simulations were done in the presence of Gα_q_ to prevent appending helices that would clash with Gα_q_ when bound to the paratope of interest. Of the 50,000 SEWING assemblies generated, the lowest energy assemblies by motif score were selected for sequence design. Each assembly was run through 1 trajectory of FastDesign^32^. During FastDesign, the backbone of Gα_q_ was fixed, but the sidechains at the interface were allowed to repack and minimize. Similarly, the backbone of the HTH at the interface on the SEWING assembly was also fixed with sidechains allowed to repack and minimize. The rest of the SEWING assembly was allowed full atom minimization and design. The 1,000 lowest energy designs were selected to undergo 10 trajectories of FastDesign, using the original SEWING assembly as the starting structure. The 10,000 designs were binned by total number of amino acids, and each bin was subjected to energy-based clustering. The 125 lowest energy cluster centers were subjected to *ab initio* structure prediction as a blind test. Of the designs with successful structure prediction and lack of prediction of alternative low energy structures, 17 were chosen for experimental validation.

For protocol 2 designs, a similar workflow was used with minor modifications. After the SEWING assembly phase, the top SEWING backbones were selected based on motif score normalized by distance of interactions in primary space. The goal was to select for backbones with higher contact order. Furthermore, the method 2 backbones were allowed to have 5 or 6 helices, whereas method 1 backbones were allowed 4 helices total. Before sequence design, the backbones were clustered based on the normalized motif score. Another filter after sequence design was a full atom relax of the design without constraints and in the absence of Gα_q_. Designs with less than 0.5 RMSD of HTH and the full SEWING assembly were selected for further analysis. The reasoning was that designs with low RMSD would be more stable and have more stable HTH motifs in the absence of Gα_q_. Prior to *ab initio* structure prediction, the lowest energy designs were filtered using a faster and lower resolution structure prediction protocol to screen 1,000 designs before subjected ∼100 to the slower more standard *ab initio* structure prediction protocol. For method 2, 22 designs were selected for experimental validation.

### Gene cloning

Genes encoding SEWN1.0 designs were ordered as clonal genes in pCDB24 from Twist bioscience. Initially, SEWN2.0 designs were planned for crystallography experiments with Gα_q/i_. Thus, the SEWN2.0 designs were cloned into the first open reading frame of pETDuet-1. The backbone of pETDuet-1 was PCR amplified using primers 5’ TAATGCTTAAGTCGAACAGAAAG 3’ and 5’ GTGGTGATGATGGTGATGGCTGC 3’. The PCR protocol was as follows: 1) 98°C for 30 sec, 2) 35 cycles of 98°C for 5 sec, 60°C for 10 sec, and 72°C for 2 min and 42 sec, and 3) 72°C for 2 min. The PCR mixture included 1X Q5 buffer, 200 μM dNTPs, 0.5 μM of each primer, 1 ng of template DNA, 1M betaine, 3% DMSO, and 0.02 U/μL Q5 polymerase. Genes encoding SEWN2.0 designs were ordered from Twist as genes with a 5’ flanking region of CATCTGAGAACCTCTACTTCCAATCA and a 3’ flanking region of TAATGCTTAAGTCGAACAGAAAGTAATC. 10 ng of amplified pETDuet-1 backbone and 10 ng gene block were incubated in Gibson assembly master mix for 2 hours at 50°C. 1μL of reaction was transformed into DH5α E. coli cells using standard heat shock protocol and then plated onto 100μg/μL ampicillin LB agar plates. Individual colonies were picked and miniprepped after overnight incubation in liquid LB at 37°C in 100μg/μL ampicillin. Clones were sequence verified before protein expression experiments.

### Protein expression and purification

SEWN designs were transformed and expressed in BL21 DE3 Star cells. Single colonies from LB agar plates supplemented with 100ug/uL ampicillin were inoculated into 5 mL of liquid LB culture. The following day, all 5 mL of culture were inoculated into 1L of TB and incubated at 37°C until OD600 = 1.0. The 1L cultures were then incubated with 1mM IPTG at 20°C for ∼20 hours before harvesting by centrifugation. Pelleted cells were resuspended in 50 mL of lysis buffer containing 100 mM HEPES pH 7.4, 300 mM NaCl, 3 mM BME, and 10 mM imidazole. Resuspended cells were passed through an Avestin C3 emulsiflex 3 times at ∼15K psi. Lysate was centrifuged at 15,000 x g for 30 minutes. Clarified lysate was run over 2 mL of pre-equilibrated Takara bio His60 Ni superflow resin. Resin was washed with 60 mL of lysis buffer with 40 mM imidazole. Purified proteins were eluted off the resin using elution buffer of 20 mM HEPES pH 7.4, 150 mM NaCl, and 250 mM imidazole.

SEWN1.0 designs were incubated with 10 ug/uL of ULP1 SUMO protease and dialyzed against 4L of 20 mM HEPES pH 7.4 and 150 mM NaCl. SEWN2.0 designs were incubated with 100 ug/uL of TEV protease and dialyzed against 4L of 20 mM HEPES pH 7.4, 150 mM NaCl, and 1 mM BME. Dialysis reactions were carried out at 4°C for ∼20 hours. Cleaved reactions were run over 2 mL of pre-equilibrated Ni resin and the flow through was collected. Collected flow through fractions were concentrated to 2.5 mL and injected onto a superdex S75 that was pre-equilibrated with 20 mM HEPES pH 7.4 and 150 mM NaCl. Purified fractions were pooled and concentrated before further experiments.

Gα_q/i_-avi in pPROEX-1 was expressed in BL21 DE3 RIPL cells. Expression and purification followed similar protocols as above with a few exceptions. All buffers were supplemented with 1 mM MgCl2, 10 uM GDP, and 1 mM BME. Following the initial Ni resin purification, Gα_q/i_ was immediately concentrated and injected onto a superdex S200 because Gα_q/i_ tends to form soluble aggregates. Afterwards, Gα_q/i_ was incubated with 100 ug/uL TEV protease overnight. Following digestion, the sample was purified the same method as above. For purification of biotinylated Gα_q/i_, after the first loading of clarified lysate onto Ni resin, the resin was washed as described and then washed again with 20 mL of biotinylation buffer 10 mM Tris HCl, 1 mM MgCl2, 10 uM GDP, and 1 mM BME. Then, the equilibrated resin was resuspended as a 50% slurry at room temperature for 1 hour in 100 ug/mL MBP-BirA enzyme, 1X avidity BiomixB, and biotinylation buffer. After 1 hour, the resin was washed with Ni washing buffer and eluted as normal.

### Circular dichroism

CD experiments were measured using a Jasco J-815 CD spectrometer. SEWN proteins in 20 mM HEPES pH 7.4 and 150 mM NaCl were diluted to 20 uM protein in CD buffer of 10 mM potassium phosphate pH 7.4 and 100 mM (NH_4_)_2_SO_4_. Samples with equivolume of SEWN buffer diluted in CD buffer were also prepared as blank controls. CD spectra were collected from 250 to 200 nm at 20°C in 1 mm quartz cuvettes using the following collection parameters: 8 second DIT, 2 nm band width, 0.5 nm data pitch, 50-100 nm/min scanning speed, and 3 accumulations. Raw CD spectra were subtracted with the corresponding blank control spectra. Thermal denaturation experiments were conducted similarly except: 1) 222 nm signal only, 2) temperature range from 2-98°C, 3) 1°C/min ramp rate, 4) and pause temperature ramp while collecting data. Guanidine hydrochloride denaturation experiments were conducted with a range of 0-6M guanidine hydrochloride with similar parameters except with a constant temperature of 20°C.

### Size exclusion chromatography with multi angle light scattering

SEC-MALS experiments were conducted on an Agilent FPLC system with a Wyatt DAWN HELEOS II light-scattering instrument, superdex 200 10/300 column, Wyatt T-rEX refractometer, and Wyatt dynamic light scattering module. The instruments were equilibrated with SEC buffer 20 mM HEPES pH 7.4,150 mM NaCl, 200 mg/L sodium azide. Protein samples were diluted to 20 uM in SEC buffer. The light scattering was collected at room temperature with a 0.5 ml/min flow rate. MALS data were analyzed using the Wyatt ASTRA software.

### Biolayer interferometry

BLI experiments were conducted using an Octet Qke instrument and Octet SA biosensors. BLI buffer consisted of 20 mM HEPES pH 7.4, 150 mM NaCl, 30 uM AlCl3, 1 mM NaF, 1 mM MgCl2, 10 uM GDP, 1 mM BME, an 1X Octet kinetics buffers. Each binding experiment consisted of 8 SA sensors being dipped into 96-well plates with the following parameters: 1) 30 sec in BLI buffer (custom), 2) 5 min in column A of fresh BLI buffer (equilibration), 3) 5 min in column B of 0.05 mg/mL Gα_q/i_-avi-biot (loading), 4) 5 min back into column A (baseline), 5) 5 min into column C of varying concentrations of SEWN protein (association), and 6) 5 min back into column A (dissociation). SEWN proteins were serially diluted from a concentration range of 0-500 nM. Raw BLI data were analyzed using ForteBio Data Analysis 9.0 with the following parameters: 1) subtraction using 0 nM SEWN trace, 2) aligning Y-axis to association, 3) inter-step correction alignment to association, 4) savitzky-golay filtering, 5) fitting association and dissociation, and 6) global fitting with Rmax unlinked. For -AlF_4_ experiments, the same experiment was conducted but with 0 mM NaF.

### Fluorescence anisotropy

FA experiments were conducted on a Horiba Jobin Yvon fluorolog-3 spectrofluorometer. FA buffer consisted of 20 mM HEPES pH 7.4, 150 mM NaCl, 30 uM AlCl3, 1 mM NaF, 1 mM MgCl2, 10 uM GDP, 1 mM BME, and 0.1% BSA. FA was measured of samples containing 0.5 uM Gα_q/i_, 10 nM FAM-HTH peptide (sequence: FAM-HQDYAEALANPIKHWSLMDQR), and SEWN protein at concentration range of 0-2 uM. The following collection parameters were used: 1) 493 excitation/517 emission, 2) polarizers on, 3) 20°C, 4) 2 nm bandwidth, 5) 1 sec integration time. FA data were fitted numerically using a python script as describe before.

### [^3^H] inositol phosphate quantitation from HEK293A cells

Low passage HEK293A cells were cultured in DMEM containing 10% FBS, 100 units/mL penicillin, and 100 ug/mL streptomycin at 37°C of 10% CO_2_. On the first day of the experiment, cells were seeded into 12-well culture dishes at a density of ∼75,000 cells per well. 24 hours later, media was aspirated and replaced with fresh media. Cells were transfected with continuum using manufacturer’s instructions and the following amounts of DNA: 1) 10 ng Gα_q_ Q209L, 2) 20 ng inhibitor plasmid, and 3) 270 ng of empty pCDNA3.1. 24 hours later, media was aspirated and replaced with Dulbecco’s inositol free media and 1uCi/well of [^3^H] inositol. 18 hours later, wells were supplemented with 10 mM LiCl for 1 hour. Afterwards, media was aspirated, and cells were lysed on ice for 30 min with 50 mM formic acid. Lysate was neutralized with 150 mM NH_4_OH and inositol phosphates were purified and quantified with Dowex chromatography.

### Luciferase activity assay

Luciferase activities were measured using the Pierce Firefly Luciferase Glow Assay Kit. Low passage HEK293A cells were maintained as described above. Cells were initially seeded and transfected as described above but with 10 ng of SRE-luciferase. After 24 hours, cells were resuspended and reseeded into 96-well plates at a density of 20,000 cells/well in serum free DMEM. After 24 hours of expression, cells were lysed and luciferase activity was measured using a clariostar plate reader as per manufacturer’s instructions.

## Supplementary Material Description

The supplementary file includes supporting experimental data, protein sequences, and additional AlphaFold2 predictions.

## Acknowledgments

This work was supported by the NIH grants R35GM131923 (B. K.) and R01GM120291(J. S.), and T32GM007040 (M. C. C.).

## abbreviations and symbols

BLI: biolayer interferometry
CD: circular dichroism
DAG: diacyl glycerol
FA: fluorescence anisotropy
Gα_q/i_: Gα_q_
G protein: guanine nucleotide-binding protein
GuHCl: guanidine hydrochloride
HTH: helix-turn-helix
IP3: inositol 1,4,5-trisphosphate
PIP2: phosphatidylinositol 4,5-bisphosphate
PLC-β: phospholipase C-β
RMSD: root mean square deviation
SEWING: structure extension with native fragment graphs

## References

1. Lombardi A, Summa CM, Geremia S, Randaccio L, Pavone V, DeGrado WF (2000) Retrostructural analysis of metalloproteins: Application to the design of a minimal model for diiron proteins. Proceedings of the National Academy of Sciences 97:6298–6305.

2. Silva D-A, Yu S, Ulge UY, Spangler JB, Jude KM, Labão-Almeida C, Ali LR, Quijano-Rubio A, Ruterbusch M, Leung I, et al. (2019) De novo design of potent and selective mimics of IL-2 and IL-15. Nature 565:186–191.

3. Sesterhenn F, Yang C, Bonet J, Cramer JT, Wen X, Wang Y, Chiang C-I, Abriata LA, Kucharska I, Castoro G, et al. (2020) De novo protein design enables the precise induction of RSV-neutralizing antibodies. Science 368.

4. Correia BE, Bates JT, Loomis RJ, Baneyx G, Carrico C, Jardine JG, Rupert P, Correnti C, Kalyuzhniy O, Vittal V, et al. (2014) Proof of principle for epitope-focused vaccine design. Nature 507:201–206.

5. Guffy SL, Pulavarti SVSRK, Harrison J, Fleming D, Szyperski T, Kuhlman B (2023) Inside-Out Design of Zinc-Binding Proteins with Non-Native Backbones. Biochemistry-us 62:770–781.

6. Azoitei ML, Correia BE, Ban YEA, Carrico C, Kalyuzhniy O, Chen L, Schroeter A, Huang PS, McLellan JS, Kwong PD, et al. (2011) Computation-Guided Backbone Grafting of a Discontinuous Motif onto a Protein Scaffold. Science 334:373–376.

7. Grigoryan G, DeGrado WF (2011) Probing Designability via a Generalized Model of Helical Bundle Geometry. J Mol Biol 405:1079–1100.

8. Huang P-S, Oberdorfer G, Xu C, Pei XY, Nannenga BL, Rogers JM, DiMaio F, Gonen T, Luisi B, Baker D (2014) High thermodynamic stability of parametrically designed helical bundles. Science 346:481–485.

9. Kuhlman B, Dantas G, Ireton GC, Varani G, Stoddard BL, Baker D (2003) Design of a Novel Globular Protein Fold with Atomic-Level Accuracy. Science 302:1364–1368.

10. Zhou J, Panaitiu AE, Grigoryan G (2020) A general-purpose protein design framework based on mining sequence–structure relationships in known protein structures. Proc National Acad Sci 117:1059–1068.

11. Harteveld Z, Bonet J, Rosset S, Yang C, Sesterhenn F, Correia BE (2022) A generic framework for hierarchical de novo protein design. Proc National Acad Sci 119:e2206111119.

12. Swanson S, Sivaraman V, Grigoryan G, Keating AE (2022) Tertiary motifs as building blocks for the design of protein-binding peptides. Protein Sci 31:e4322.

13. Wang J, Lisanza S, Juergens D, Tischer D, Watson JL, Castro KM, Ragotte R, Saragovi A, Milles LF, Baek M, et al. (2022) Scaffolding protein functional sites using deep learning. Science 377:387–394.

14. Watson JL, Juergens D, Bennett NR, Trippe BL, Yim J, Eisenach HE, Ahern W, Borst AJ, Ragotte RJ, Milles LF, et al. (2022) Broadly applicable and accurate protein design by integrating structure prediction networks and diffusion generative models.

15. Ingraham J, Baranov M, Costello Z, Frappier V, Ismail A, Tie S, Wang W, Xue V, Obermeyer F, Beam A, et al. (2022) Illuminating protein space with a programmable generative model.

16. Torres SV, Leung PJY, Lutz ID, Venkatesh P, Watson JL, Hink F, Huynh H-H, Yeh AH- W, Juergens D, Bennett NR, et al. (2022) De novo design of high-affinity protein binders to bioactive helical peptides. Biorxiv:2022.12.10.519862.

17. Lipsh-Sokolik R, Khersonsky O, Schröder SP, Boer C de, Hoch S-Y, Davies GJ, Overkleeft HS, Fleishman SJ (2023) Combinatorial assembly and design of enzymes. Science 379:195–201.

18. Jacobs TM, Williams B, Williams T, Xu X, Eletsky A, Federizon JF, Szyperski T, Kuhlman B (2016) Design of structurally distinct proteins using strategies inspired by evolution. Science [Internet] 352:687–690. Available from: http://www.sciencemag.org/cgi/doi/10.1126/science.aad8036

19. Guffy SL, Teets FD, Langlois MI, Kuhlman B (2018) Protocols for Requirement-Driven Protein Design in the Rosetta Modeling Program. J Chem Inf Model 58:895–901.

20. Gresset A, Sondek J, Harden TK (2012) Phosphoinositides I: Enzymes of Synthesis and Degradation. Subcell Biochem 58:61–94.

21. Lyon AM, Tesmer JJG (2013) Structural Insights into Phospholipase C-β Function. Mol Pharmacol 84:488–500.

22. Kamato D, Mitra P, Davis F, Osman N, Chaplin R, Cabot PJ, Afroz R, Thomas W, Zheng W, Kaur H, et al. (2017) Gaq proteins: molecular pharmacology and therapeutic potential. Cell Mol Life Sci 74:1379–1390.

23. Raamsdonk CDV, Bezrookove V, Green G, Bauer J, Gaugler L, O’Brien JM, Simpson EM, Barsh GS, Bastian BC (2008) Frequent somatic mutations of GNAQ in uveal melanoma and blue naevi. 457:599–602.

24. D. VRC, G. GK, B. CM, C. GM, Swapna V, Thomas W, C. OA, Werner W, Gary G, Nancy B, et al. (2010) Mutations in GNA11 in Uveal Melanoma. New Engl J Med 363:2191– 2199.

25. Kawakami K, Yanagawa M, Hiratsuka S, Yoshida M, Ono Y, Hiroshima M, Ueda M, Aoki J, Sako Y, Inoue A (2022) Heterotrimeric Gq proteins act as a switch for GRK5/6 selectivity underlying β-arrestin transducer bias. Nat Commun 13:487.

26. Nagai J, Bellafard A, Qu Z, Yu X, Ollivier M, Gangwani MR, Diaz-Castro B, Coppola G, Schumacher SM, Golshani P, et al. (2021) Specific and behaviorally consequential astrocyte Gq GPCR signaling attenuation in vivo with iβARK. Neuron 109:2256–2274.e9.

27. Cabezudo S, Sanz-Flores M, Caballero A, Tasset I, Rebollo E, Diaz A, Aragay AM, Cuervo AM, Mayor F, Ribas C (2021) Gαq activation modulates autophagy by promoting mTORC1 signaling. Nat Commun 12:4540.

28. Schrage R, Schmitz A-L, Gaffal E, Annala S, Kehraus S, Wenzel D, Büllesbach KM, Bald T, Inoue A, Shinjo Y, et al. (2015) The experimental power of FR900359 to study Gq-regulated biological processes. Nat Commun 6:10156.

29. Zhang H, Nielsen AL, Strømgaard K (2020) Recent achievements in developing selective Gq inhibitors. Med. Res. Rev. 40:135–157.

30. Waldo GL, Waldo GL, Ricks TK, Ricks TK, Hicks SN, Hicks SN, Cheever ML, Cheever ML, Kawano T, Kawano T, et al. (2010) Kinetic Scaffolding Mediated by a Phospholipase C- and Gq Signaling Complex. Science 330:974–980.

31. Hussain M, Cummins MC, Endo-Streeter S, Sondek J, Kuhlman B (2021) Designer proteins that competitively inhibit Gαq by targeting its effector site. J Biological Chem 297:101348.

32. Maguire JB, Haddox HK, Strickland D, Halabiya SF, Coventry B, Griffin JR, Pulavarti SVSRK, Cummins M, Thieker DF, Klavins E, et al. (2021) Perturbing the energy landscape for improved packing during computational protein design. Proteins Struct Funct Bioinform 89:436–449.

33. Charpentier TH, Waldo GL, Lowery-Gionta EG, Krajewski K, Strahl BD, Kash TL, Harden TK, Sondek J (2016) Potent and Selective Peptide-based Inhibition of the G Protein Gαq. The Journal of biological chemistry 291:25608–25616.

34. Robinson-Rechavi M, Godzik A (2005) Structural Genomics of Thermotoga maritima Proteins Shows that Contact Order Is a Major Determinant of Protein Thermostability. Structure 13:857–860.

35. Yarwood RE, Imlach WL, Lieu T, Veldhuis NA, Jensen DD, Herenbrink CK, Aurelio L, Cai Z, Christie MJ, Poole DP, et al. (2017) Endosomal signaling of the receptor for calcitonin gene-related peptide mediates pain transmission. Proc National Acad Sci 114:12309–12314.

36. Gorvin CM, Rogers A, Hastoy B, Tarasov AI, Frost M, Sposini S, Inoue A, Whyte MP, Rorsman P, Hanyaloglu AC, et al. (2018) AP2σ Mutations Impair Calcium-Sensing Receptor Trafficking and Signaling, and Show an Endosomal Pathway to Spatially Direct G-Protein Selectivity. Cell Reports 22:1054–1066.

37. Jensen DD, Lieu T, Halls ML, Veldhuis NA, Imlach WL, Mai QN, Poole DP, Quach T, Aurelio L, Conner J, et al. (2017) Neurokinin 1 receptor signaling in endosomes mediates sustained nociception and is a viable therapeutic target for prolonged pain relief. Sci Transl Med 9.

38. Jimenez-Vargas NN, Pattison LA, Zhao P, Lieu T, Latorre R, Jensen DD, Castro J, Aurelio L, Le GT, Flynn B, et al. (2018) Protease-activated receptor-2 in endosomes signals persistent pain of irritable bowel syndrome. Proc National Acad Sci 115:E7438–E7447.

