## supplementary_materials for "*De novo* design of stable proteins that efficaciously inhibit oncogenic G proteins"

**running title**: Rosetta de novo design of $G\alpha_q$ inhibitors

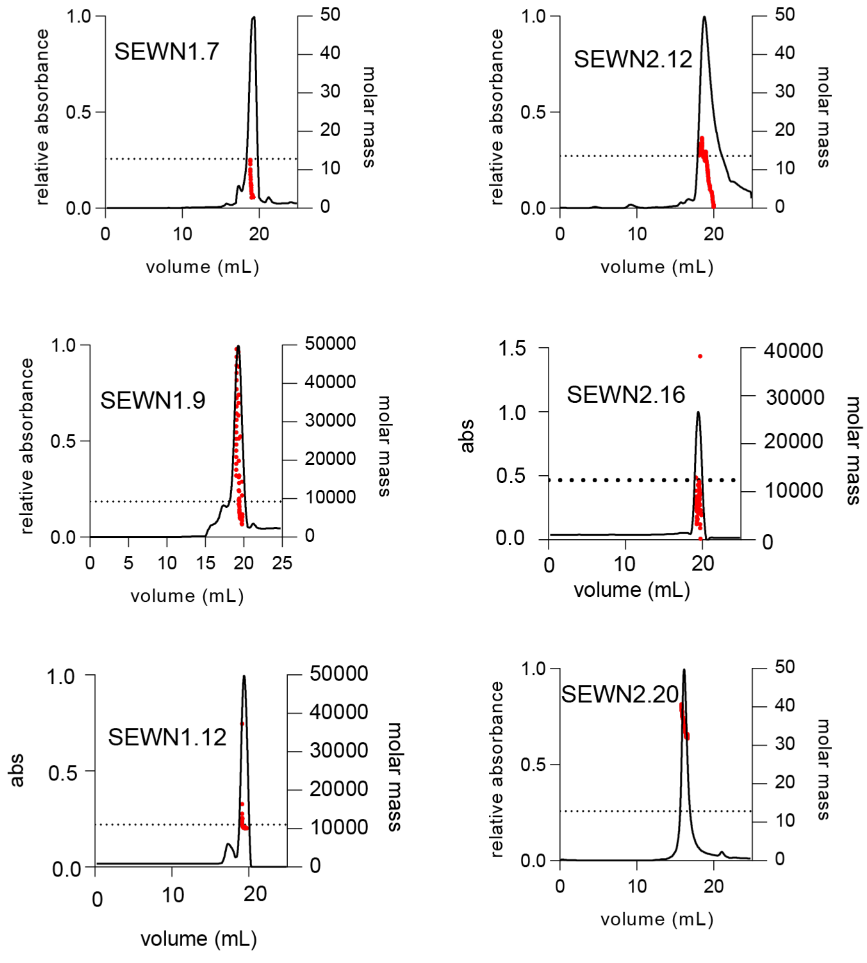

**Figure S1 SEC-MALS of SEWN proteins**

All SEWN proteins were run at a concentration of 20 μM on a Superdex S200 size exclusion column. Dotted lines represent theoretical molecular weight of a monomer. Except for SEWN2.20, the SEWN proteins elute near the bed volume of 24mL. Thus, the light scattering was noisy and precise molecular weights were difficult to obtain. However, SEWN2.20 clearly elutes before the buffer with a strong light scattering with a molecular weight corresponding to a tetramer.

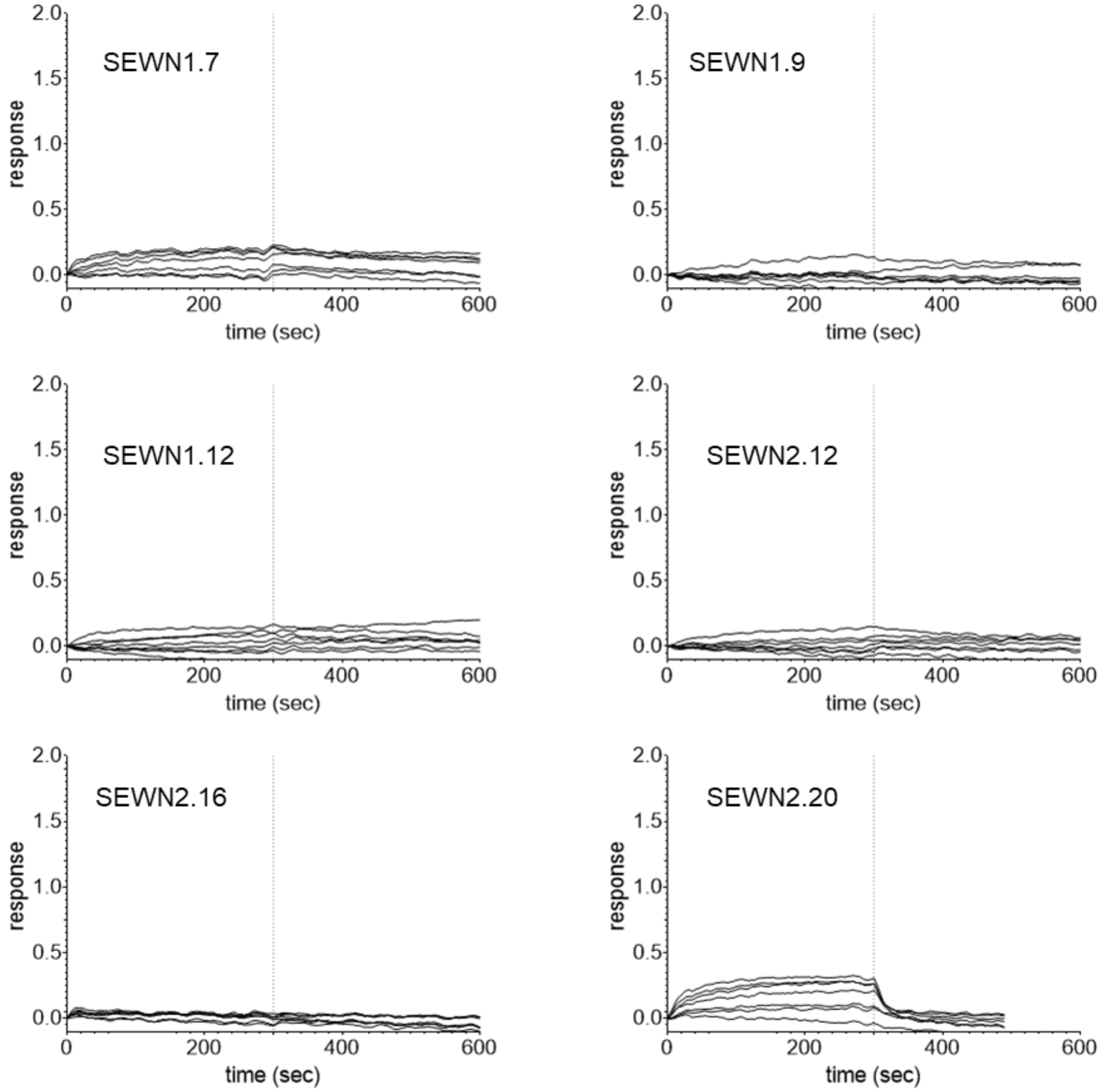

**Figure S2 SEWN proteins do not bind the inactive form of G$\alpha_q$**
The same BLI experiment was performed as in Figure 3, with the exception that $AlF_4^-$ was excluded from the buffer and the highest [SEWN] is 1 μM as opposed to 500 nM. Plots are scaled to the same axis as in Figure 3. The SEWN proteins show more than 15-fold loss of response.

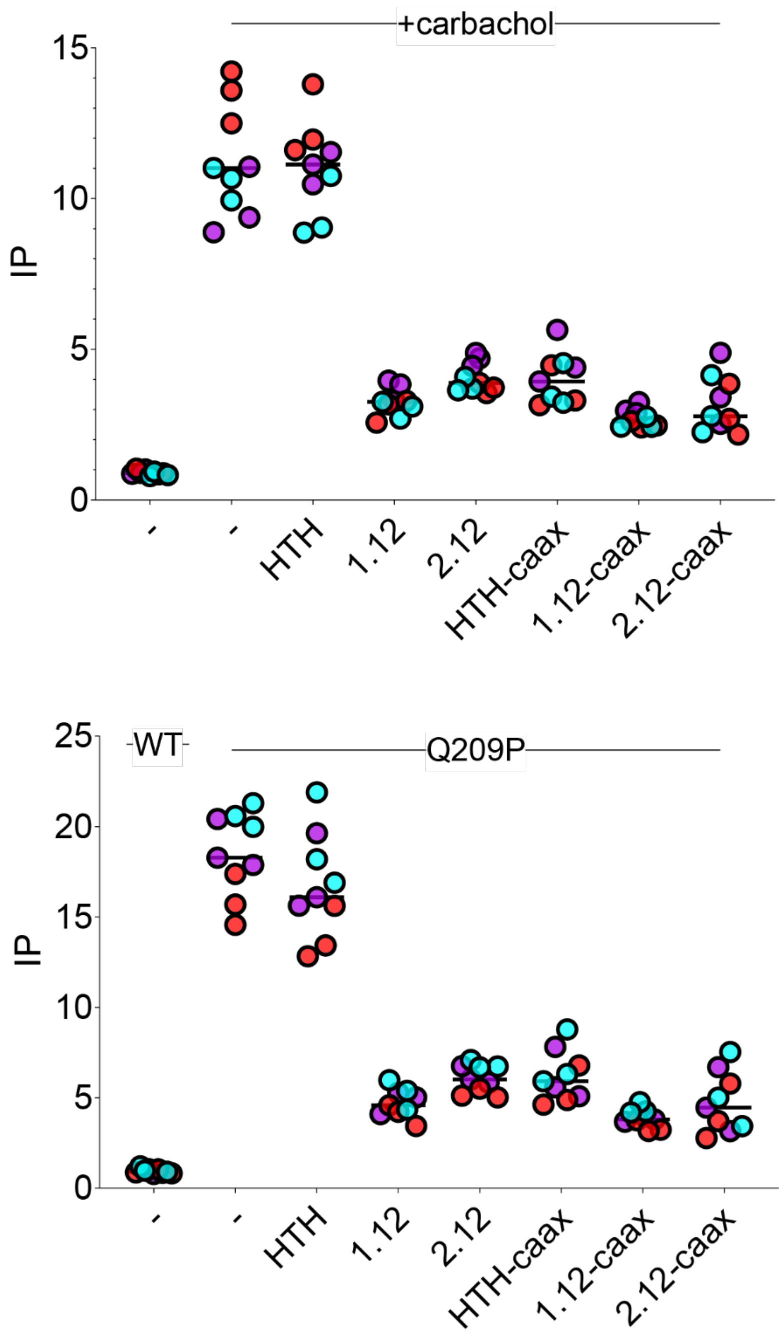

**Figure S3 SEWN1.12 and 2.12 inhibit G$\alpha_q$ WT and Q209P in HEK293A cells**

G$\alpha_q$ activation of PLC isozymes in HEK293A was triggered using either carbachol or transfecting G$\alpha_q$ Q209P. Simultaneously, different inhibitors were co-transfected into the HEK293A cells. Across both experiments, the SEWN proteins with or without caax inhibit G$\alpha_q$ Q209P with the same efficacy as HTH-caax.

MSYYHHHHHHDYDIPTTENLYFQGAMGCTLSAEDKAAVERSKMIDRNLREDGEKAAR
EVKLLLLGTGESGKSTFIKQMRIIHGSGYSDEDKRGFTKLVYQNIFTAMQAMIRAMDTL
KIPYKYEHNKAHAQLVREVDVEKVSAFENPYVDAIKSLWNDPGIQECYDRRREYQLSD
STKYYLNDLDRVADPSYLPTQQDVLRVRVPTTGIIEYPFTFKDLHFKMVDVGGQRSER
RKWIHCFEGVTAIIFCVALSDYDQVLVESDNENRMEESKALFRTIITYPWFQNSSVILFL
NKKDLLEEKIMYSHLVDYFPEYDGPQRDAQAAREFILKMFVDLNPDSDKIIYSHFTCAT
DTENIRFVFAAVKDTILQLNLKEYNLVGLNDIFEAQKIEWHE
His-G$\alpha_{q/i}$-avi

MGHHHHHHHHHHGSLQDSEVNQEAKPEVKPEVKPETHINLKVSDGSSEIFFKIKKTTP
LRRLMEAFAKRQGKEMDSLRFLYDGIRIQADQAPEDLDMEDNDIIEAHREQIGGGSEIM
KTLEKLQKKIKEMRKNNDEESVRKLETLVYAMALINPIVYWLAMDLERIDRLLKEMEKN
NDEDSVKEMRERKERLRKMIEEAKKS
HisSUMO-SEWN1.7

MGHHHHHHHHHHGSLQDSEVNQEAKPEVKPEVKPETHINLKVSDGSSEIFFKIKKTTP
LRRLMEAFAKRQGKEMDSLRFLYDGIRIQADQAPEDLDMEDNDIIEAHREQIGGSLEE
MQELLKALEKDPNNPEIKTKTYAKALINPIVYWLLMDARWVREALKTGGHPDEREKLKE
QLKKVETKLKEIQKKD
HisSUMO-SEWN1.9

MGHHHHHHHHHHGSLQDSEVNQEAKPEVKPEVKPETHINLKVSDGSSEIFFKIKKTTP
LRRLMEAFAKRQGKEMDSLRFLYDGIRIQADQAPEDLDMEDNDIIEAHREQIGGSEAD
EIASKLEERVRRVAKTLEEMTKQGDEDSTEKMVRLLIKLEEEIKRIAKKYNNPPKMQKLL
KTLQTEMYAMALINPIIWWRVMDKYKKS
HisSUMO-SEWN1.12

MGSSHHHHHHGSSENLYFQSTDEAIHEVAKMLRLDEEIIKRLIQKDDRAREALKRFEEIL
QRLRKAEKNGDEELQRRLRKILQQLLKELAQLIYAMALINPIEVWRLLEQAQHK
His-SEWN2.12

MGSSHHHHHHGSSENLYFQSDDQVERMIHEEGLDVEQIIEILIRRIRDPKLQKALKDAR
ELLQLQKLLEELKRQGASEEELKQIRKRIHQLALINPIWWWMAVQIRRKQ
His-SEWN2.16

MGSSHHHHHHGSSENLYFQSDDELLKEVERILHEIAKHGDKRLIEHAMRLLQQIEQLIQ
KKKKDGVSDEELKELIKKLHQLALINPIVWWLIILARQMNVDLQQALQIL
His-SEWN2.20

5FAM-HQDYAEALANPIKHWSLMDQR
FAM-HTH

MEQKLISEEDLGSGSGSSEADEIASKLEERVRRVAKTLEEMTKQGDEDSTEKMVRLLIK
LEEEIKRIAKKYNNPPKMQKLLKTLQTEMYAMALINPIIWWRVMDKYKKS
Myc-SEWN1.12

MEQKLISEEDLGSGSGSSEADEIASKLEERVRRVAKTLEEMTKQGDEDSTEKMVRLLIK
LEEEIKRIAKKYNNPPKMQKLLKTLQTEMYAMALINPIIWWRVMDKYKKSKLAAAQLYT
RASQPELAPEDPEDLEPPPVKKRKRKCAIL
Myc-SEWN1.12-Rac1caax

MEQKLISEEDLGSGSGSTDEAIHEVAKMLRLDEEIIKRLIQKDDRAREALKRFEEILQRL
RKAEKNGDEELQRRLRKILQQLLKELAQLIYAMALINPIEVWRLLEQAQHK
Myc-SEWN2.12

MEQKLISEEDLGSGSGSTDEAIHEVAKMLRLDEEIIKRLIQKDDRAREALKRFEEILQRL
RKAEKNGDEELQRRLRKILQQLLKELAQLIYAMALINPIEVWRLLEQAQHKKLAAAQLYT
RASQPELAPEDPEDLEPPPVKKRKRKCAIL
Myc-SEWN2.12-Rac1caax

**Figure S4 sequences of proteins used in this study**
All His-tagged sequences were expressed in E. coli. All Myc-tagged sequences were expressed in HEK293 cells. The FAM-tagged HTH was used in fluorescence anisotropy experiments only.

**Table S1 Table of fitted rates of SEWN designs binding biotinylated G$\alpha_q$**

| design | $K_D$ (M) | $k_{on}$ (M/sec) | $k_{off}$ (1/sec) |
|---|---|---|---|
| 1.7 | $<1.0\times10^{-9}$ | $3.9\times10^{4}\pm 3.5\times10^{3}$ | $<1.0\times10^{-7}$ |
| 1.9 | $5.3\times10^{-8}\pm 1.4\times10^{-9}$ | $7.8\times10^{4}\pm 9.0\times10^{2}$ | $4.1\times10^{-3}\pm 2.5\times10^{-5}$ |
| 1.12 | $4.9\times10^{-8}\pm 8.5\times10^{-9}$ | $1.5\times10^{4}\pm 3.7\times10^{2}$ | $7.2\times10^{-4}\pm 1.9\times10^{-5}$ |
| 2.12 | $2.0\times10^{-8}\pm 6.3\times10^{-10}$ | $9.0\times10^{4}\pm 1.0\times10^{3}$ | $1.8\times10^{-3}\pm 1.8\times10^{-5}$ |
| 2.16 | $6.3\times10^{-8}\pm 5.0\times10^{-9}$ | $1.5\times10^{4}\pm 2.6\times10^{2}$ | $9.5\times10^{-4}\pm 1.3\times10^{-5}$ |
| 2.20 | $8.3\times10^{-8}\pm 6.8\times10^{-9}$ | $9.9\times10^{3}\pm 2.2\times10^{2}$ | $8.2\times10^{-4}\pm 1.1\times10^{-5}$ |

### Table S2 AlphaFold2 predictions of SEWN1.12 and SEWN2.12 agree well with Rosetta models.

AlphaFold2 predictions were made with the sequence of G$\alpha_q$ included in the query sequence. The displayed RMSD is for all C$\alpha$ atoms in the design model.

| design | RMSD after superimposing G$\alpha_q$ chain (Å) | RMSD after superimposing the design chains (Å) |
|---|---|---|
| 1.7 | 2.6 | 2.6 |
| 1.9 | 4.4 | 1.5 |
| 1.12 | 2.2 | 0.9 |
| 2.12 | 2.1 | 1.8 |
| 2.15 | 27.6 | 1.7 |
| 2.20 | 20.5 | 6.9 |
